# Predicting Antibody Developability from Sequence using Machine Learning

**DOI:** 10.1101/2020.06.18.159798

**Authors:** Xingyao Chen, Thomas Dougherty, Chan Hong, Rachel Schibler, Yi Cong Zhao, Reza Sadeghi, Naim Matasci, Yi-Chieh Wu, Ian Kerman

**Author notes:** Equal contribution (Xingyao Chen, Thomas Dougherty, Chan Hong, Rachel Schibler, Yi Cong Zhao). Correspondence to: Yi-Chieh Wu < >.

## Abstract

Antibodies are prominent therapeutic agents but costly to develop. Existing approaches to predict developability depend on structure, which requires expensive laboratory or computational work to obtain. To address this issue, we present a machine learning pipeline to predict developability from sequence alone using physicochemical and learned embedding features. Our approach achieves high sensitivity and specificity on a dataset of 2400 antibodies. These results suggest that sequence is predictive of developability, enabling more efficient development of antibodies.

## 1. Introduction

Since the United States Food and Drug Administration approved the first monoclonal antibody (mAb) in 1986, therapeutic antibodies have exploded in popularity due to their high specificity and few adverse effects (Lu et al., 2020), and now have a global market value of US $115.2 billion (Lu et al., 2020). However, there are significant barriers to manufacturing mAbs at an industrial scale, and bringing a therapeutic antibody to market can cost US $1.4 billion and take up to 12 years (Mestre-Ferrandiz et al., 2012).

To be clinically effective, mAbs must be present in high concentrations (Chames et al., 2009). Therefore, candidates for therapeutic use must retain specificity and safety throughout development, while maintaining high stability and low aggregation to be fit for industrial production. A strategy to minimize failure is to exclude candidates early in the development cycle based on their aggregation propensity. One such metric, Developability Index (DI), relies on an antibody’s hydrophobic and electrostatic interactions as inferred from its three-dimensional structure (Lauer et al., 2012).

However, researchers often do not have structural data for newly proposed antibodies. Experimental approaches to determine structure are expensive, costing up to US $100,000 per protein (Yang et al., 2018b), and computational protein structure prediction through homology modeling (Lauer et al., 2012) or deep learning (Senior et al., 2020) is error-prone and time-consuming. These limitations make estimating developability difficult for large collections of candidates or to explore large numbers of potential variants.

Machine learning approaches have successfully replaced the need for structure in several protein prediction tasks (Long et al., 2018; Rahman et al., 2016). These approaches often require only the sequence as input and are thus more widely applicable than methods that rely on structure.

In this work, we present a machine learning pipeline to predict DI directly from sequence, thereby bypassing the need to determine structure experimentally or computationally. With our sequence-based method, researchers can screen candidate sequences for therapeutic antibodies in bulk, identifying those with high potential for industrial development and production. To validate our approach, we applied it to a dataset of 2400 antibodies with known sequence and structure. Our pipeline achieved high sensitivity and specificity, indicating that sequence is predictive of developability.

## 2. Methods

We framed our problem as a supervised machine learning task, with sequence as input and developability index as output. In this section, we present the main steps in our pipeline (Figure 1).

**Figure 1.**
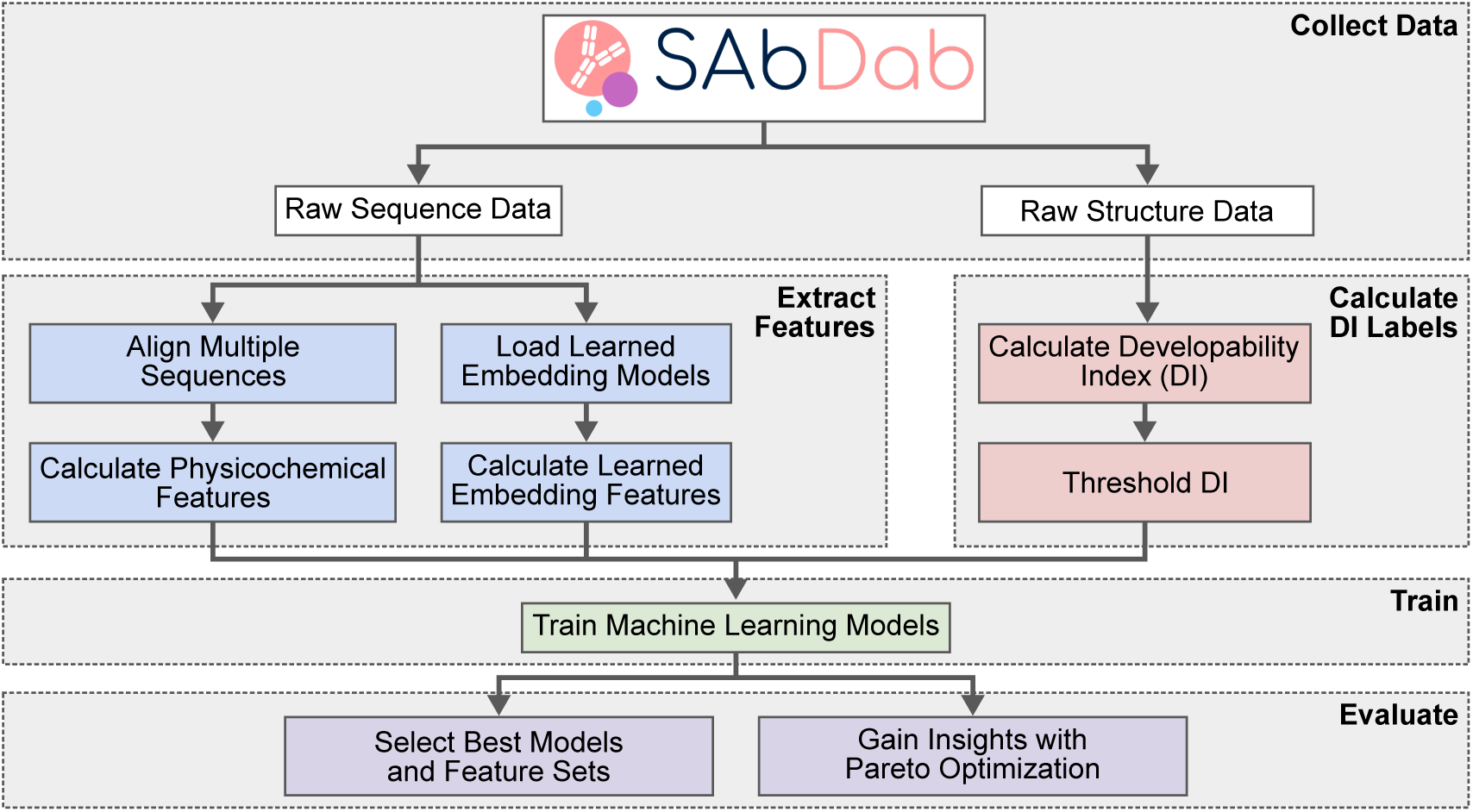
Machine learning pipeline. After collecting data from SAbDab (white), for each antibody, we extract feature vectors from sequence (blue) and calculate developability index labels from structure (red). We use these together to train several machine learning models (green), tuning hyperparameters through cross-validation (not shown). Lastly, we evaluate our approach using both standard machine learning metrics and Pareto optimization (purple). [SAbDab logo from http://opig.stats.ox.ac.uk/webapps/newsabdab/sabdab/.]

### 2.1. Datasets

We gathered antibody data from the Structural Antibody Database (SAbDab, Dunbar et al., 2014). From an initial dataset of 3816 antibodies, we retained 2426 antibodies that satisfy the following criteria:

1. have both sequence (FASTA) and Protein Data Bank (PDB) structure files,
2. contain both a heavy chain and a light chain, and
3. have crystal structures with resolution < 3 Å (Wlodawer et al., 2007).

As only the variable regions of the heavy and light chains are used to compute DI, we extracted exactly one heavy and one light chain. We also removed extraneous protein chains and heteroatoms from the PDB structure files to ensure that the calculated DI reflects only an antibody’s variable region.

### 2.2. Label Generation

We calculated the antibody DI values using ‘Calculate DI’, part of BIOVIA’s Pipeline Pilot (Dassault Systémes BIOVIA, 2020). We were unable to calculate DI values for 17 antibodies because of the presence of non-standard residues and other unknown errors. Though DI values are continuous, we decided to frame our task as a classification problem rather than regression, as classification is an easier prediction task and it is more robust to noisy data. As a low DI value corresponds to high developability, we thresholded DI values, with the bottom 20% as developable and the top 80% as non-developable. Though an 80-20 split creates an imbalanced dataset, our goal is to create a pool enriched in candidates with a high chance of being developable. We treated the binary DI labels as the “ground truth” for our supervised learning models. Our final dataset contains 2409 antibodies sequences with binary DI labels.

### 2.3. Feature Extraction and Preprocessing

We extracted two broad types of features for each antibody sequence: (1) physicochemical properties derived directly from sequence, and (2) vectors in an embedded space similar to the popular doc2vec (Le & Mikolov, 2014) model.

#### 2.3.1. Physicochemical Features

First, we computed a simple feature that measures the percentage of each amino acid in the sequence.

Then, based on our knowledge of DI (Lauer et al., 2012), we selected physicochemical features expected to be relevant. These features include 9 whole sequence-based properties (isoelectric point, molecular weight, average residue weight, charge, molar extinction coefficient, molar extinction coefficient of cystine bridge, extinction coefficient, extinction coefficient of cystine bridge, and improbability of expression in inclusion bodies (IEIB)), computed with EMBOSS (Rice et al., 2000) and Expasy (Gasteiger et al., 2005).

In addition, we computed several physicochemical features based on amino acid-based properties (Kyte-Doolittle hydropathy, hydrophobic moment, and charge). However, using amino acid-based properties presents a challenge because our machine learning models require fixed-length feature vectors as inputs but the sequence lengths of antibodies can vary. Therefore, we explored multiple approaches to obtain fixed-length vectors: (1) exclude the amino acid-based features; (2) pad amino acid-based features with zero; (3) pad amino acid-based features with the feature’s average value in the sequence; (4) replace amino acid-based features with summary statistics (mean, median, and standard deviation); and (5) align amino acid-based features using a multiple sequence alignment (MSA). In the last strategy, we align the sequences using ClustalW (Thompson et al., 1994), align amino acid-based features based on the MSA, then impute gaps by using the average feature value across all sequences for that position in the MSA.

Lastly, we standardized the features by removing the mean and scaling to unit variance. In total, we explored six different physicochemical feature sets.

#### 2.3.2. Learned Embedding Features

Learned embedding features are based on the word2vec (Mikolov et al., 2013) and doc2vec (Le & Mikolov, 2014) models, which produce embeddings by mapping words and documents to vectors of real numbers. Similar word embeddings have been proposed for *n*-grams in biological sequences, for example, BioVec, ProtVec, and GeneVec (Asgari & Mofrad, 2015). Such embeddings can infer biological properties of unseen sequences without requiring an understanding of the underlying physical or biological mechanisms.

In this work, we used the embedding models presented in Yang et al., 2018a. There, the authors divided protein sequences into non-overlapping *k*-mers (1 ≤ *k* ≤ 5), learned embeddings that place *k*-mers that occur in similar contexts near each other, then considered multiple *k*-mers in a fixed window size. We input our antibody sequences into these pre-trained embedding models to vectorize each sequence. In total, we explored 149 embedding feature sets.

### 2.4. Experimental Setup

Using `scikit-learn` (Pedregosa et al., 2011), we evaluated one baseline (that generates predictions by respecting the training set’s class distribution) and six (real) models (Table 1). To train and evaluate our models, we used an 80-20 stratified train-test split. To tune hyperparameters, we used 10-fold stratified cross-validation. For each model, we randomly sampled hyperparameters and selected the best set of hyperparameters based on the mean validation F_1_ score.

**Table 1.** Machine learning models and hyperparameters.

| model | hyperparameters |
| --- | --- |
| Stratified Sampling | – |
| Gaussian Bayes | – |
| Logistic Regression | regularization $\in \{L2\}$<br>$C \in \text{loguniform}(0.001, 1000)$ |
| Support Vector Machine | $C \in \text{loguniform}(0.001, 100)$<br>kernel $\in \{\text{linear}, \text{rbf}\}$<br>$\gamma \in \text{loguniform}(0.001, 1)$ |
| Random Forest | # of estimators $\in \{1, 10, \dots, 200\}$<br>max depth $\in \{0, 2, \dots, 50\}$<br>max fraction features $\in \{0.1, 0.15, \dots, 0.75\}$ |
| Gradient Boosting | learning rate $\in \text{loguniform}(0.01, 0.5)$<br># of estimators $\in \{1, 10, \dots, 200\}$<br>max depth $\in \{0, 2, \dots, 20\}$<br>max fraction features $\in \{0.1, 0.2, \dots, 0.6\}$ |
| Multilayer Perceptron | hidden layer sizes<br>$\in \{(100, ), (50, ), (100, 100)\}$ |

We evaluated model performance using several standard machine learning metrics, including Area Under the Receiver Operating Characteristics (AUROC) curve, Area Under the Precision-Recall (AUPR) curve, F_1_ score, precision, and recall. For deployment, we suggest the best model-feature pair based on *F*_1_ score.

However, because our results may depend on the metric used, we also applied Pareto optimization to select models and feature sets on the Pareto front.^1^ Such models and feature sets are optimal in the sense that one metric cannot be increased without decreasing at least one other metric. Prior to Pareto optimization, we filtered our models to ensure they meet a baseline, requiring performance under each metric of at least 0.4.

## 3. Results

### 3.1. Performance across Features and Models

To determine how performance varies across model-feature set pairs, we generated a heatmap of mean validation *F*_1_ scores (Figure 2). Unsurprisingly, using similar feature sets tends to yield similar performances regardless of model. Furthermore, features based on embedding models with 1-mers perform poorly (mean *F*_1_ < 0.32). For every other feature set, every machine learning model outperformed our baseline model (mean *F*_1_: 0.36−0.57), with Support Vector Machine (mean *F*_1_: 0.52 − 0.65) and Multilayer Perceptron (mean *F*_1_: 0.50 − 0.64) performing best. Of the feature sets, physicochemical (mean *F*_1_ for all non-baseline models: 0.51 − 0.60) and learned embeddings with *k*-mers of sizes 2 − 4 (mean *F*_1_: 0.50 − 0.54) perform best.

**Figure 2.**
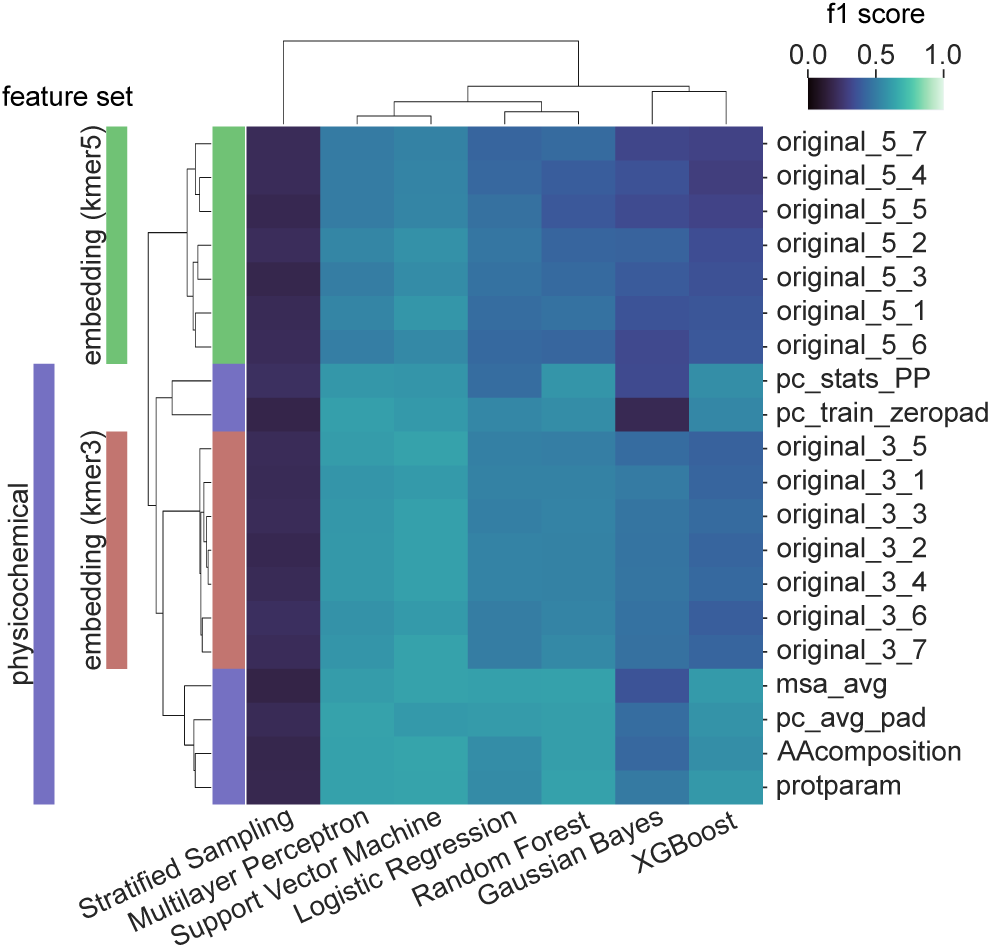
Average validation *F*_1_ scores of all models trained on a selection of feature sets. Models and feature sets are clustered by score similarity. Of the models, Support Vector Machine and Multilayer Perceptron perform best. Of the feature sets, physicochemical and embedding feature sets perform similarly. The exception is that embedding feature sets using *k*-mers of size *k* = 1 perform very poorly (not shown). Using *k*-mers of size *k* = 2 and *k* = 4 perform similarly to *k* = 3 (not shown). A description of the feature sets can be found in Table S1.

Given that physicochemical and embeddings feature sets perform similarly, we would prefer to use physicochemical features as they are more easily interpretable. Overall, the best combination of model type and feature set was the Support Vector Machine trained on physicochemical features with multiple sequence alignment.

### 3.2. Performance of Various Models using the Best Feature Set

Next, we used the best feature set (physicochemical features with multiple sequence alignment) and investigated the performance of the various models (Figure S1). Though Gaussian Bayes performs poorly, all other non-baseline models achieve high training performance. However, these models also generalize poorly, indicating overfitting.

### 3.3. Performance of Various Feature Sets using the Best Model

Similarly, we used the best model (Support Vector Machine) and investigated the performance of the various feature sets (Figure S2). The top two physicochemical feature sets and the top two embedding feature sets show similar performance, and again, there is evidence of overfitting.

### 3.4. Model and Feature Selection by Pareto Optimality

Finally, rather than optimizing on only one metric, *F*_1_ score, we looked at the Pareto front, which simultaneously considers five metrics (AUPR, AUROC, *F*_1_, precision, and recall). A model or feature set that occurs frequently in the Pareto front indicates that it performs well under several metrics. The Pareto front contains 148 combinations of model - feature - hyperparameter set. Feature sets that appear the most frequently are physicochemical features and embedding features with 3-mers. Model types that appear the most frequently are Support Vector Machine and Random Forest. Importantly, these results are consistent with our analysis of performance based solely on *F*_1_ score.

## 4. Discussion

In this manuscript, we have presented a machine learning pipeline that extracts features derived from antibody sequence data to predict its developability. By using only sequence-based features, we remove the need to experimentally determine or computationally predict antibody structures.

While our results demonstrate that an antibody’s developability index is predictable using machine learning, we must be wary of using DI as a measure of an antibody’s true potential. Because DI is computationally determined based on aggregation propensity, it may ignore other indicators of developability. Further investigation using a curated database is needed to determine how well aggregation propensity correlates with actual developability.

Furthermore, our analysis of performance is based on a relatively small dataset of antibodies. This limitation resulted in our models overfitting to the training data and generalizing poorly. Future work could augment the dataset with simulations that introduce point mutations into sequences and computationally predict the associated structure. While this approach would introduce artifacts from structural prediction, it would also enable more complex regression or deep learning models. Recent work has shown that deep learning models using convolutional neural networks or long short-term memory models can predict protein function from primary sequence (Bileschi et al., 2019; Kulmanov & Hoehndorf, 2020).

As with any machine learning approach, our pipeline is flexible and can be retrained on a larger dataset with experimentally-validated labels to potentially achieve a model with greater accuracy and predictive power.

## Acknowledgements

The authors thank Zachary Dodds, Kathy Ryan, and Surani Gunasena at Harvey Mudd College for help with Harvey Mudd College Clinic program, and Lisa Yan at Dassault Systèmes BIOVIA for help developing the project outline.

This work was supported by the Department of Computer Science of Harvey Mudd College and by Dassault Systèmes BIOVIA.

---

**Table S1.** Feature sets.

| feature | description |
| --- | --- |
| AAcomposition | percentage of each amino acid in sequence |
| pc_stats_PP | summary statistics of amino acid-based physicochemical features with sequence-based physicochemical features provided by EMBOSS |
| protparam | amino-acid composition and sequence-based physicochemical features provided by ExPASy Proteomics Server (Gasteiger et al., 2005) |
| pc_train_zeropad | sequence- and amino acid-based physicochemical features provided by EMBOSS, with zero padding |
| pc_avg_pad | sequence- and amino acid-based physicochemical features provided by EMBOSS, with average padding |
| msa_avg | sequence- and amino acid-based physicochemical features provided by EMBOSS, with aligned sequences and gaps imputed using average value |
| original_X_Y | embedding features with $k$-mer size X ($1 \leq X \leq 5$) and window size Y ($1 \leq Y \leq 7$) |

**Figure S1.**
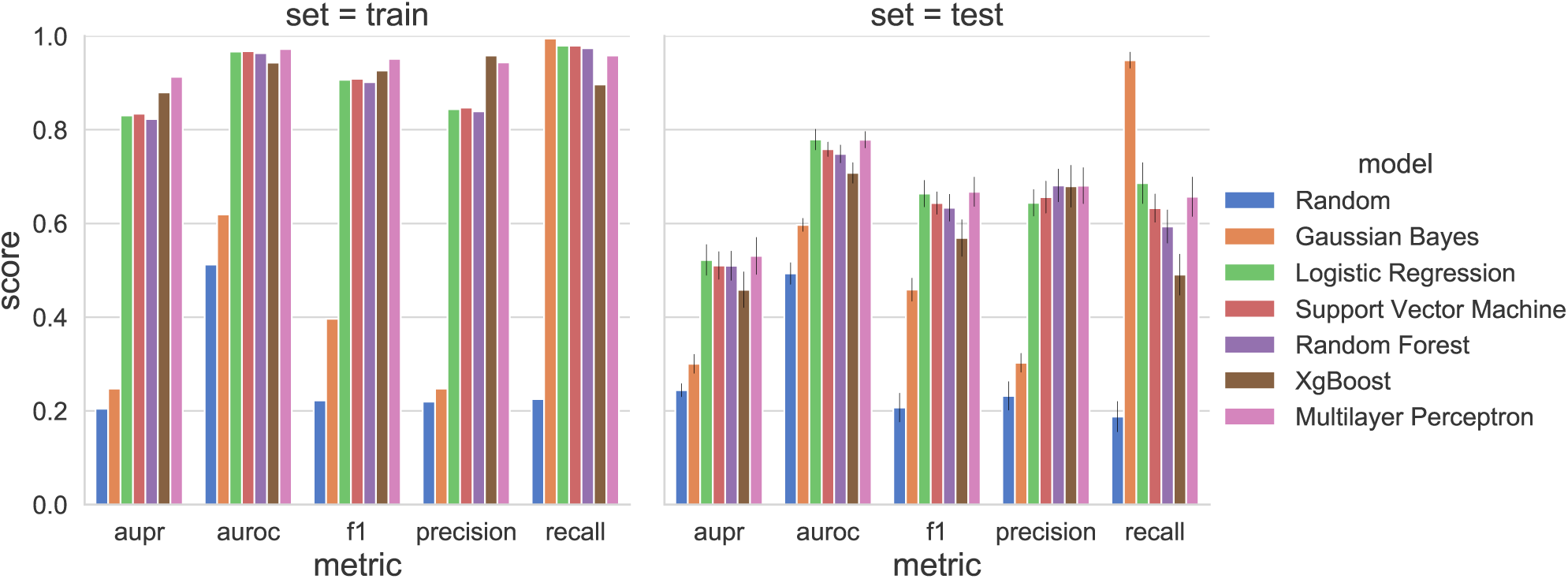
Training set (left) and test set (right) performance of all models trained on the top physicochemical feature set. Error bars represent the standard deviation across 10 bootstraps.

**Figure S2.**
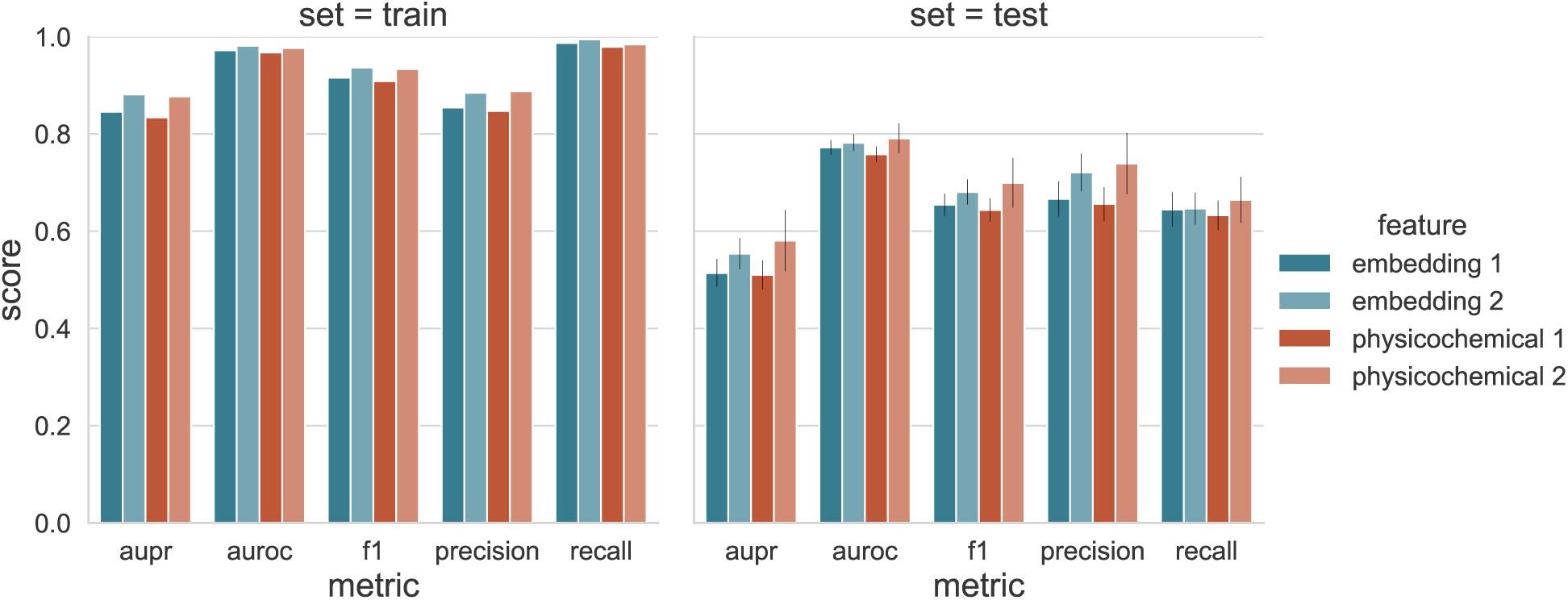
Training set (left) and test set (right) performance of Support Vector Machines trained on the top two embedding features (original_3_7 and original_3_5) and the top two physicochemical features (msa_avg and protparam).

## Footnotes

1 Formally, we measure the performance of each model-feature set pair using a vector of scores *v* = ⟨AUPR, AUROC, *F*_1_, precision, recall⟩. Given two vectors *v* and *v*′, *v* is said to be *strictly better* than *v*′ if each entry of *v* is greater than or equal to the corresponding entry in *v*′ and at least one entry of *v* is greater than its corresponding entry in *v*′. Given a set *V* of vectors, *v* ∈*V* said to be *Pareto-optimal* with respect to *V* if there does not exist any other *v*′ ∈*V* that is strictly better than *v*. The *Pareto front* is the set of vectors that are all Pareto-optimal.

